# Tension suppresses Aurora B kinase-triggered release of reconstituted kinetochore-microtubule attachments

**DOI:** 10.1101/415992

**Authors:** Anna K. de Regt, Cordell J. Clark, Charles L. Asbury, Sue Biggins

## Abstract

Chromosome segregation requires kinetochores to attach to microtubules from opposite spindle poles. Proper attachments come under tension and are stabilized, but defective attachments lacking tension are released, giving another chance for correct attachments to form. This error correction process requires the Aurora B kinase, which phosphorylates kinetochores to destabilize microtubule attachments. However, the mechanism by which Aurora B can distinguish kinetochore tension remains unclear because it is difficult to detect kinase-triggered detachment and manipulate kinetochore tension *in vivo*. To address these challenges, we developed an optical trapping-based flow assay with soluble Aurora B and reconstituted kinetochore-microtubule attachments. Strikingly, we found that tension on these attachments suppressed their Aurora B-triggered release, suggesting that tension-dependent changes in the conformation of kinetochores can regulate Aurora B activity or its outcome. Our work uncovers the basis for a key mechano-regulatory event that ensures accurate segregation and may inform studies of other mechanically regulated enzymes.

## Introduction

It has been known for decades that tension signals the proper attachment of chromosomes to spindle microtubules and causes selective stabilization of these attachments (Nicklas and Koch, 1969; Nicklas and Ward, 1994), but how tension suppresses the destabilizing activity of Aurora B remains unclear (Biggins et al., 1999; Cheeseman et al., 2006; Cimini et al., 2006; DeLuca et al., 2006; Dewar et al., 2004; Lampson et al., 2004; Mukherjee et al., 2019; Tanaka et al., 2002). A popular explanation has been that tension-dependent stretching of chromosomes might spatially separate the prominent inner-centromeric pool of Aurora B from kinetochores, thereby inhibiting their phosphorylation (Liu et al., 2009; Tanaka et al., 2002). However, the inner-centromeric localization of Aurora B is dispensable for accurate chromosome segregation and additional kinetochore binding sites for the kinase have recently been uncovered (Broad et al., 2020; Campbell and Desai, 2013; Fischbock-Halwachs et al., 2019; Hengeveld et al., 2017; Ruchaud et al., 2007; Yue et al., 2008), suggesting that other mechanisms might underlie error correction. An alternate ‘substrate conformation’ model proposes that the structural changes kinetochores undergo when they come under tension block Aurora B phosphorylation (Aravamudhan et al., 2014; Campbell and Desai, 2013; Hengeveld et al., 2017; Ruchaud et al., 2007; Smith et al., 2016; Uchida et al., 2009; Wan et al., 2009). Electron microscopy, fluorescence microscopy, and biochemical crosslinking studies have demonstrated that kinetochores undergo large structural changes when they come under tension, which could potentially block Aurora B phosphorylation via substrate occlusion or another mechanism (Joglekar et al., 2009; Tien et al., 2014; Wan et al., 2009). This idea is similar to known instances where tension-dependent conformational changes regulate kinase enzymes in other cellular contexts (Bauer et al., 2019; Baumann et al., 2017; Puchner et al., 2008; Sawada et al., 2006).

It has not been possible to rigorously test models for Aurora B regulation within cells for several reasons. Aurora B has additional mitotic roles, so inhibition of the kinase acts globally and not specifically on error correction (Broad and DeLuca, 2020; Willems et al., 2018). Moreover, it is challenging to test various models *in vivo* because unattached kinetochores also lack tension, and kinetochores that make attachments quickly come under tension. Finally, the phosphorylation state of individual kinetochores *in vivo* cannot be precisely correlated with kinase activity because substrate phosphorylation represents a balance of both kinase and opposing phosphatase activity (Foley et al., 2011; Kruse et al., 2013; Liu et al., 2010; Suijkerbuijk et al., 2012). To overcome these challenges, we developed a reconstituted system that allows direct observation of Aurora B-triggered detachments at different levels of tension using optical trapping. Strikingly, we found that tension on kinetochores inhibits soluble Aurora B from detaching kinetochores from microtubules, demonstrating that tension can directly suppress the outcome of Aurora B activity.

## Results

### Optical trap flow assay to test Aurora B-triggered release of kinetochore-microtubule attachments

To test the substrate conformation model, we sought to establish an assay in which individual, reconstituted kinetochore-microtubule attachments can be placed under high or low tension using a laser trap, subsequently exposed to soluble active Aurora B kinase, and then monitored until detachment (Fig. 1A). Our previous reconstitution using native kinetochores isolated from budding yeast served as a foundation (Akiyoshi et al., 2010). We also required a source of purified, soluble Aurora B kinase with sufficient activity to quickly phosphorylate its substrates before a kinetochore would typically detach from a microtubule. Aurora B is a member of the <u>C</u>hromosomal <u>P</u>assenger <u>C</u>omplex (CPC) and is activated by the C-terminus of the CPC component INCENP (Krenn and Musacchio, 2015).

**Figure 1.**
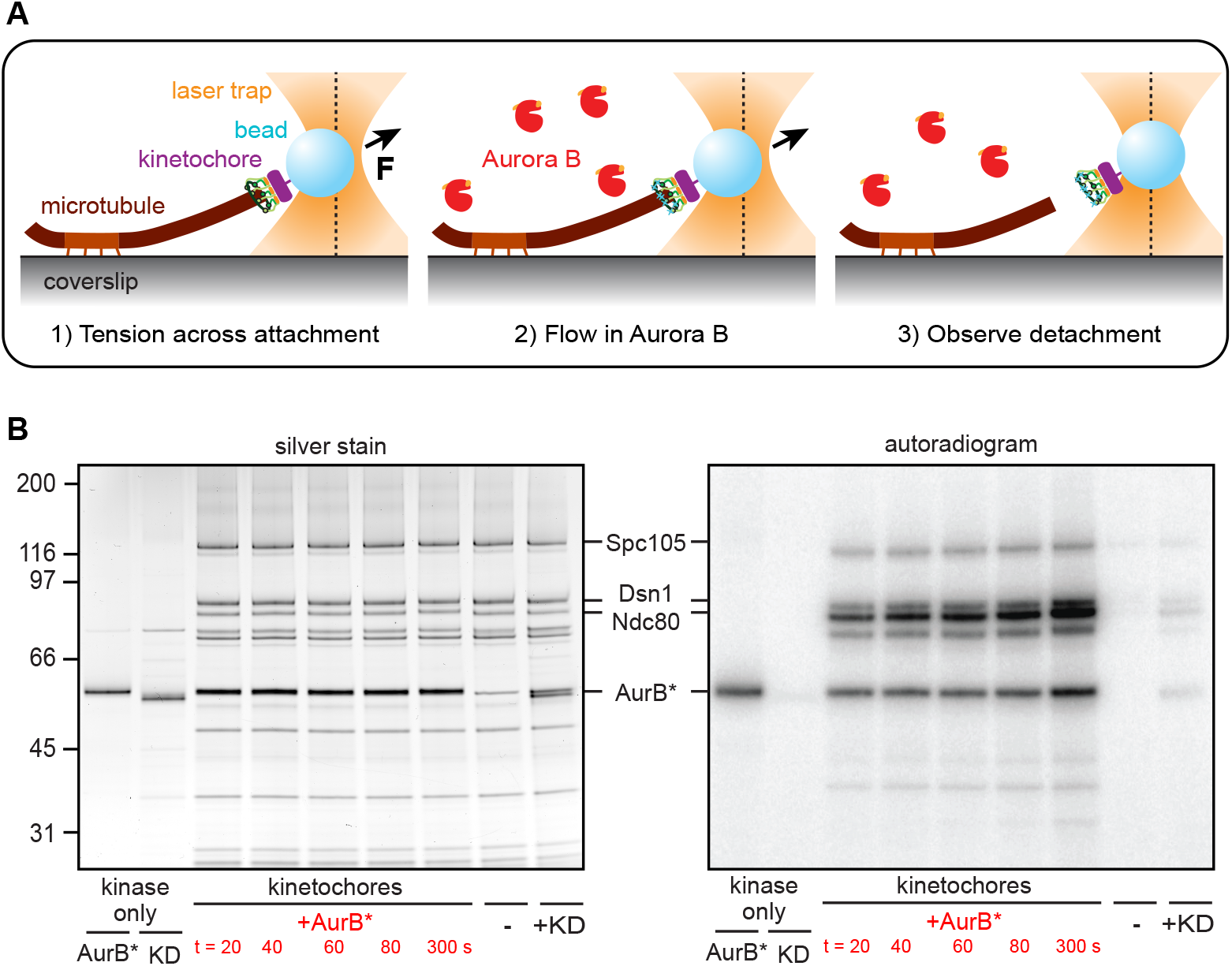
Optical trap flow assay for testing whether tension affects Aurora B-triggered release of kinetochore-microtubule attachments. (**A**) Schematic of the assay. A kinetochore-decorated bead is attached to the growing tip of a microtubule anchored to a coverslip. Tension is applied with an optical trap, and free kinase is introduced by gentle buffer exchange. Tip-tracking of the kinetochore-bead is monitored until detachment or interruption of the experiment. (**B**) An engineered Aurora B construct phosphorylates kinetochores rapidly. Mps1-1 kinetochores (purified from SBY8726) were incubated with either 0.2 μM AurB* or 0.2 μM AurB*-KD in the presence of 32P-γ-labeled ATP and then visualized by silver stain and autoradiography. The first two lanes show AurB* or AurB*-KD alone with no kinetochores after 5 minutes of incubation, the next five lanes show a time course (seconds, s) of kinetochores incubated with AurB*, and each of the last two lanes show kinetochores incubated with either no kinase or AurB*-KD for 3.5 minutes. The relevant Aurora B substrates are labeled and the positions of protein standards (kDa) are shown on the left. Ndc80 incorporation of 32P is quantified in Figure S1B.

We made a highly active recombinant Aurora B kinase lacking the inner centromere targeting domains by fusing the yeast INCENP^Sli15^ activation box (residues 624-698) to the N-terminus of yeast Aurora B^Ipl1^, creating a construct we call AurB*. We confirmed its activity by incubating purified kinetochores with ATP-γ-^32^P and either AurB* or a kinase-dead mutant (AurB*-KD), in which the catalytic lysine was replaced with arginine (Fig. 1B). Kinetochores purified from wild-type cells lack endogenous Aurora B kinase, and also the inner centromere, but they co-purify with endogenous Mps1 kinase (Akiyoshi et al., 2010; London et al., 2012). Therefore, we used kinetochores from a temperature-sensitive mutant *mps1-1* strain that had been shifted to the non-permissive temperature to inactivate Mps1. Consistent with previous findings (Meyer et al., 2013), the Mps1-1 kinetochores lacked co-purifying Dam1 complex (Dam1c; Fig. S1A), but otherwise resemble wild-type purified kinetochores (London et al., 2012). In addition, the kinetochores were purified in the presence of irreversible phosphatase inhibitors so we could directly monitor the effects of Aurora B independently of any opposing phosphatase activity. We found that AurB* autoactivated (via autophosphorylation) and efficiently phosphorylated known kinetochore substrates Ndc80, Spc105 and Dsn1 (London et al., 2012) (Fig. 1B). The key microtubule-coupling substrate Ndc80 was phosphorylated to greater than 50% completion in under one minute with 0.2 μM AurB* (Figs. 1B and S1B). In contrast, kinetochores incubated with AurB*-KD or only ATP-γ-^32^P exhibited little phosphorylation, confirming the lack of endogenous kinase activity. These experiments show that AurB* can rapidly phosphorylate kinetochores.

### Aurora B phosphorylates physiological substrates to decrease kinetochore-microtubule affinity

To test whether AurB* phosphorylates physiological targets, we repeated the ATP-γ-^32^P experiment using phospho-deficient mutant kinetochores with alanine substitutions at the seven known Aurora B target sites on the Ndc80 N-terminal ‘tail’ (Ndc80-7A) (Akiyoshi et al., 2009). Because combining the *ndc80-7A* and *mps1-1* mutations in yeast is lethal, we used kinetochores isolated from strains containing wild-type Mps1 for this experiment. Mps1 also phosphorylates Ndc80 (Sarangapani et al., 2021), so Ndc80 was partially phosphorylated even in the absence of exogenous AurB*, along with other kinetochore proteins (Fig 2A). However, the addition of AurB* increased phosphorylation on wild-type kinetochores compared to mutant Ndc80-7A kinetochores (Fig. 2A). Thus, AurB* phosphorylates kinetochores on physiologically-relevant sites.

**Figure 2.**
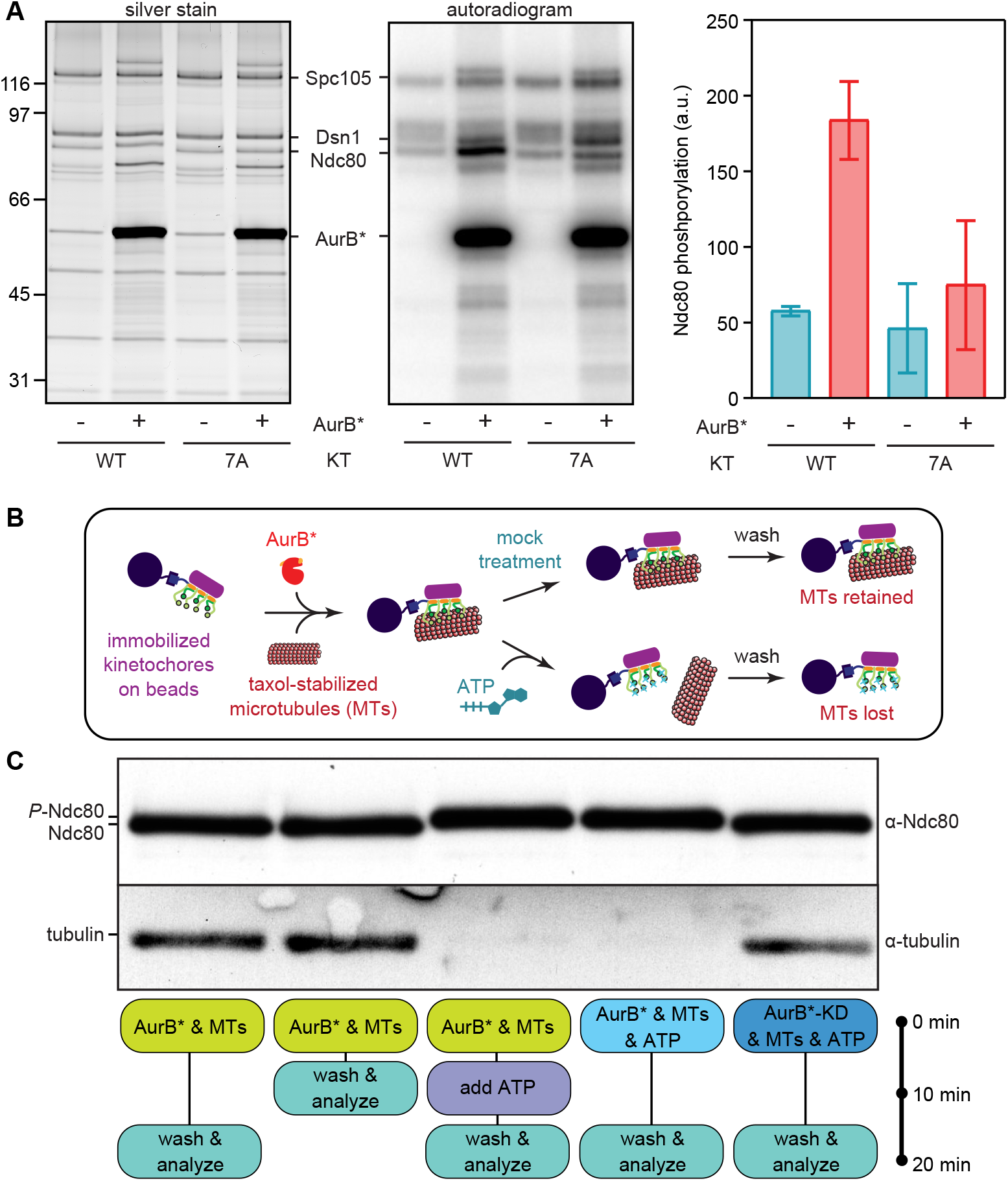
Aurora B phosphorylates physiological substrates and reduces kinetochore-microtubule affinity. (**A**) Wild type (SBY8253) or phospho-deficient mutant Ndc80-7A (SBY8522) kinetochores (KTs) were incubated with either buffer or 1 μM AurB* in the presence of 32P-γ-labeled ATP for 3 minutes. A decrease in phosphorylation of the Ndc80 band when comparing lanes 2 and 4 shows that AurB* phosphorylates Ndc80 on one or more of the seven alanine-substituted residues in the phospho-deficient mutant. Ndc80 phosphorylation is quantified at right. (**B**) Schematic for the experiment shown in (C). (**C**) Purified Mps1-1 kinetochores (SBY8726) immobilized on beads were mixed at the indicated times with taxol-stabilized microtubules (MTs), ATP, and AurB* or AurB*-KD, and then washed and analyzed. Components remaining bound to the beads were separated by SDS-PAGE and analyzed by immunoblot. *P*-Ndc80: phosphorylated Ndc80.

To test whether AurB* is sufficient to reduce kinetochore-microtubule affinity, we purified Mps1-1 kinetochores, retained them on the purification beads, and then incubated the beads with taxol-stabilized microtubules together with either AurB* or AurB*-KD. We subsequently added buffer or ATP, washed the beads, and quantified the bound microtubules by immunoblotting (Fig. 2B). As expected, kinetochores bound to microtubules in the absence of ATP (Fig. 2C, first and second lanes). When ATP was added, there was an AurB*-dependent release of the microtubules (Fig. 2C, lanes 3 through 5) that occurred more slowly if phosphorylation of Ndc80 was blocked at the Aurora B phospho-sites (Fig. S2) (Akiyoshi et al., 2009). These observations show that phosphorylation by AurB* is sufficient to release taxol-stabilized microtubules from kinetochores and confirm that one or more of the known phospho-sites contributes to this release.

### Aurora B activity is sufficient to release kinetochores from dynamic microtubule tips

To determine whether Aurora B can release kinetochores from dynamic microtubule tips, which are the physiological substrate for error correction, we linked Mps1-1 kinetochores to polystyrene microbeads and then used an optical trap to attach the beads individually to the assembling tips of single, coverslip-anchored microtubules (Akiyoshi et al., 2010; Miller et al., 2016; Sarangapani et al., 2013). We included native purified Dam1c in the assay buffer to ensure the Mps1-1 kinetochores could remain persistently attached to growing and shortening tips under a variety of tensile forces (Fig. S3) (Akiyoshi et al., 2010; Gutierrez et al., 2020). We adapted the trapping assay to introduce AurB* by flowing kinase-containing buffer into the reaction chamber after a kinetochore-microtubule tip attachment had been established and placed under tension (Figs. 1A and 3A). This approach was technically challenging. Unlike previous versions of our optical trap assay, only one kinetochore attachment could be probed per slide, because any free-floating unattached kinetochores became phosphorylated once AurB* was introduced and thus were unsuitable for further measurements. Moreover, flow in the chamber sometimes caused extra beads or aggregated proteins to catch in the laser, ending the events prematurely (Fig. S4A). We therefore made a number of modifications to the assay in order to improve data collection efficiency, including increasing the density of kinetochores on the beads, which avoided lengthy searches for active beads. We applied a constant, relatively low tension of ~1 pN between a kinetochore-coated bead and a microtubule (Akiyoshi et al., 2010), introduced AurB*-KD or AurB* at one of two different concentrations, and then monitored the bead until either detachment or interruption. Tip-tracking of the kinetochore-beads was usually undisturbed by the onset of flow (Figs. 3A, S4A, and S4B). The spontaneous detachment rate when the control AurB*-KD was flowed in was 2.3 ± 0.9 hr^−1^ and this was elevated with active AurB* in a concentration-dependent manner, either two-fold to 4.6 ± 1.4 hr^−1^ (0.5 μM AurB*) or three-fold to 7.2 ± 2.2 hr^−1^ (5 μM AurB*) (Figs. 3B, S4C and S4D). Analyzing the detachment rates specifically from shortening versus growing microtubule tips showed that the introduction of active AurB* led to more frequent detachment regardless of the tip state (Figs. 3C and 3D). Together, these data show that AurB* activity is sufficient to detach kinetochores from dynamic microtubule tips that are either growing or shortening *in vitro*.

**Figure 3.**
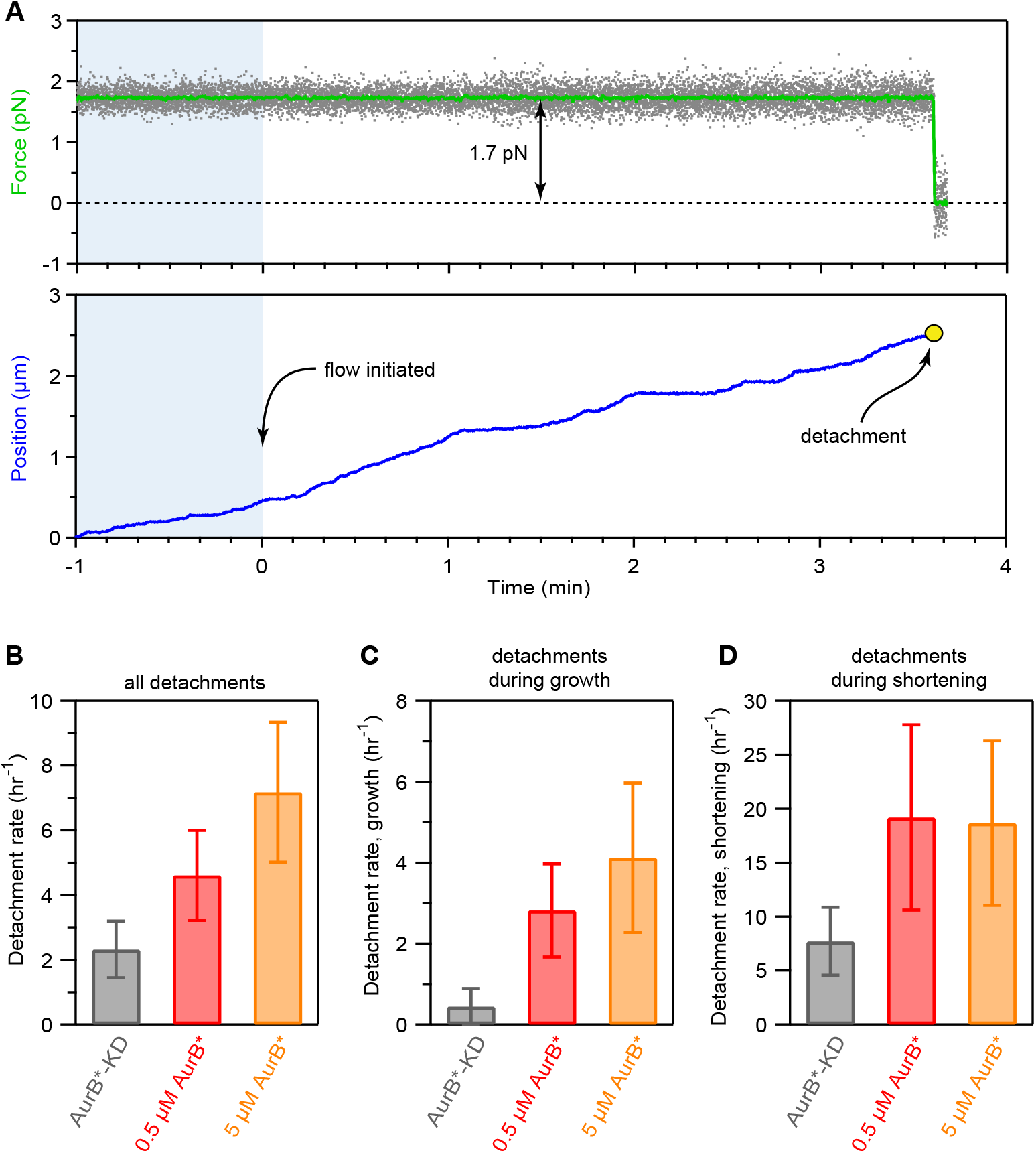
Aurora B activity is sufficient to release kinetochores supporting low tension from the tips of dynamic microtubules. (**A**) Example record of optical trap data for an individual detachment event. Gray points show the force applied and green trace shows the same data after smoothing. Blue trace shows the relative position of the kinetochore-decorated bead over time. Kinase introduction occurred at time 0; blue shading indicates data recorded before kinase introduction. See Figure 1A for a schematic of the experiment. (**B**) Overall detachment rates for kinetochore-decorated beads supporting ~1 pN of tension in the presence of 5 μM AurB*-KD, 0.5 μM AurB*, or 5 μM AurB*. (**C**, **D**) Detachment rates measured specifically during microtubule growth (C) or shortening (D) for the same conditions as in (B). Data for (B – D) were collected using a high density of kinetochores on the trapping beads (Dsn1:bead ratio, 3,300). Error bars represent uncertainty due to Poisson statistics. Numbers of detachments, rate estimates, and statistical comparisons are detailed in Supplementary Data Table 1.

### Tension suppresses Aurora B-triggered detachment

To test the effect of tension on Aurora B-triggered detachment, we sought to repeat the AurB* flow experiment at two different levels of laser trap force, 1.5 and 5 pN, chosen to span the estimated range of physiological forces on yeast kinetochores *in vivo* (Chacon et al., 2014; Mukherjee et al., 2019). During our initial low-force experiments that used beads densely decorated with Mps1-1 kinetochores, spontaneous detachments from shortening microtubules were unusually rare compared to our previous work (Fig. S5A), which used beads decorated much more sparsely with wild-type kinetochores (Akiyoshi et al., 2010; Miller et al., 2016; Sarangapani et al., 2013). Lowering the density of Mps1-1 kinetochores on the beads restored the higher detachment rates (Fig. S5B), suggesting that at high densities, multiple kinetochores could share the applied load (Fig. S5C). Because such load-sharing will reduce the amount of tension per kinetochore, potentially to a level insufficient to inhibit Aurora B, we reverted to a sparse decoration of kinetochores on the beads for all subsequent experiments. Active beads were more difficult to find, as expected, but the Mps1-1 kinetochores in the presence of Dam1c showed a catch bond-like increase in attachment stability with force, consistent with our previous studies using wild-type kinetochores (Akiyoshi et al., 2010; Miller et al., 2016) (Fig. S5D). To measure the effect of tension on Aurora B-triggered release, we established kinetochore-microtubule attachments at either low (1.5 pN) or high tension (5 pN), flowed in either AurB*-KD or AurB*, and monitored the detachment rates. AurB* activity strongly promoted kinetochore release at low force, elevating the detachment rate from 10.4 ± 3.3 hr^−1^ in kinase-dead controls up to 28.9 ± 5.3 hr^−1^ when active AurB* was introduced (Fig. 4), a roughly three-fold increase similar to what we observed earlier using densely decorated beads at low force. However, at high force, the detachment rates in the presence of AurB* versus AurB*-KD were statistically indistinguishable, and both around 7 hr^−1^. These observations show that tension on the reconstituted kinetochore-microtubule attachments suppressed their release by AurB*.

**Figure 4.**
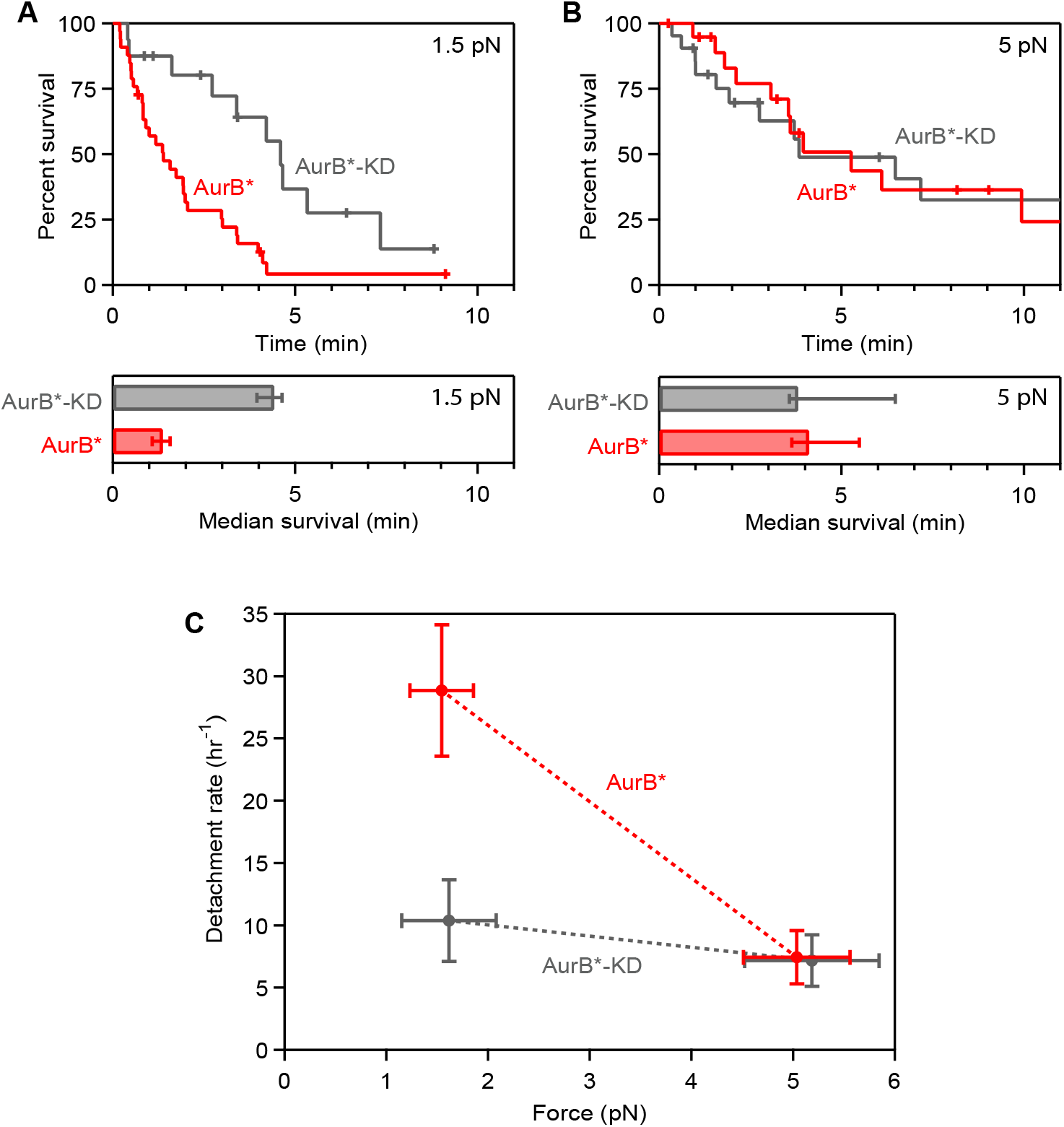
Tension suppresses Aurora B-triggered detachment. (**A**, **B**) Kaplan-Meier survival analysis (upper plots) and median survival times (lower bar graphs) for kinetochore-microtubule attachments supporting either ~1.5 pN (A) or ~5 pN of tension (B) in the presence of 0.5 μM AurB* or 0.5 μM AurB*-KD. Tick marks on Kaplan-Meier plots represent censored data from interrupted events that did not end in detachment. Based on log-rank tests, p-values for the data shown in (A) and (B) are 0.0016 and 0.98, respectively (kinase-dead versus active AurB*). Median survival bar graphs show times at which the estimated Kaplan-Meier survival probability falls below 50%. Error bars represent interquartile range, estimated by bootstrapping. (**C**) Overall detachment rates for kinetochore-decorated beads in the presence of 0.5 μM AurB* or 0.5 μM AurB*-KD as a function of the applied force. All data for (A – C) were collected using a low density of kinetochores on the trapping beads (Dsn1:bead ratio, 200). Vertical error bars in (C) represent uncertainty due to Poisson statistics. Horizontal error bars in (C) represent standard deviation. Numbers of detachments, rate estimates, and statistical comparisons are detailed in Supplementary Data Table 1.

## Discussion

A number of models have been proposed to explain how Aurora B selectively weakens kinetochore-microtubule attachments that lack tension, while leaving load-bearing, tip-coupled kinetochores unmodified (Aravamudhan et al., 2014; Campbell and Desai, 2013; Hengeveld et al., 2017; Liu et al., 2009; Ruchaud et al., 2007; Smith et al., 2016; Tanaka et al., 2002; Uchida et al., 2009; Wan et al., 2009). Rigorously testing such models requires independent control of tension, attachment, and enzyme activity. Here, we developed an assay that fulfills these needs and allows components to be introduced while kinetochore-microtubule attachments are held continuously under tension. While previous work has shown that pre-phosphorylation reduces the microtubule binding affinity of individual kinetochore components (Cheeseman et al., 2006; Sarangapani et al., 2013; Umbreit et al., 2012), our results provide the first direct demonstration that Aurora B activity is sufficient to detach kinetochores from dynamic microtubule tips, consistent with recent work suggesting that Aurora B’s inner centromere localization is dispensable for error correction in cells (Campbell and Desai, 2013; Hengeveld et al., 2017; Yue et al., 2008). The flow assay we developed here can be used in the future to test how tension affects other enzymes involved in error correction.

Because our experiments used soluble AurB*, the tension we applied was not directly borne by the kinase. Therefore, the tension must have altered the kinetochores or the microtubule tips in a way that prevented soluble AurB* from triggering detachments. This substrate conformation effect can explain how relaxed kinetochore-microtubule attachments are selectively released *in vivo* while tension-bearing attachments persist (Dewar et al., 2004; Mukherjee et al., 2019). Tension causes kinetochores to undergo large structural changes *in vivo* (Dumont et al., 2012; Joglekar et al., 2009; Maresca and Salmon, 2009; Uchida et al., 2021; Wan et al., 2009), and such changes could alter the ability of Aurora B to trigger kinetochore-microtubule detachments. We speculate that the key Aurora B substrate residues might become masked when a kinetochore comes under tension, or perhaps the substrates are inhibited from threading into the kinase active site (Fig. 5). It is also formally possible that Aurora B phosphorylation is not affected, per se, and that the tension in our experiments instead suppressed the effects of phosphorylation. In any case, the underlying mechanism must differ from that of the other mechanically regulated kinases uncovered to date, which are all activated rather than inhibited by tension (Bauer et al., 2019; Baumann et al., 2017; Puchner et al., 2008; Sawada et al., 2006).

**Figure 5.**
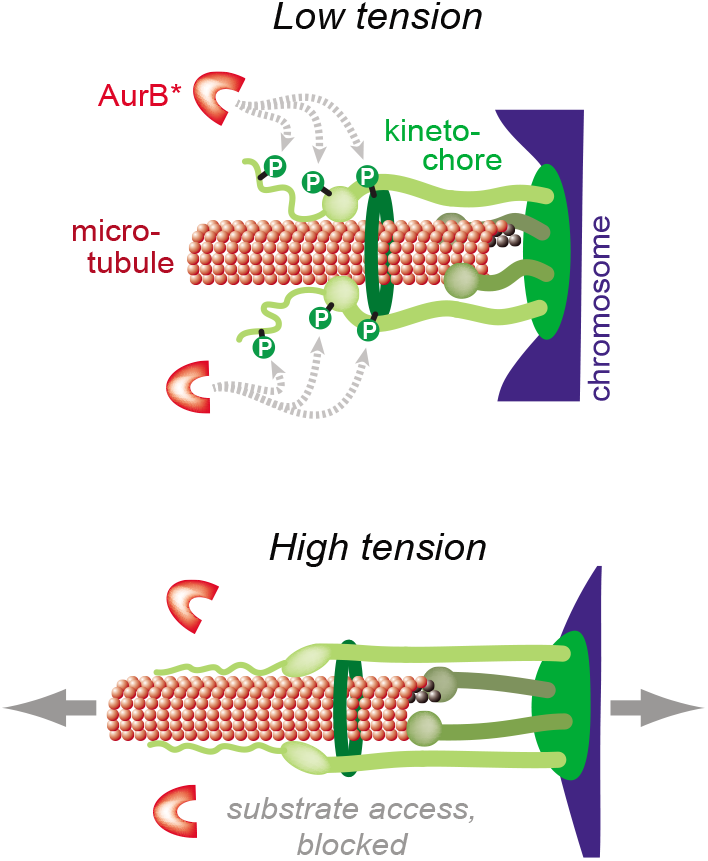
Model illustrating how conformational changes within a kinetochore could inhibit Aurora B-triggered detachment. The key Aurora B substrate residues might become masked when a kinetochore comes under tension, or the substrates could be inhibited from threading into the kinase active site.

Mechano-regulation of Aurora B via the substrate conformation mechanism does not exclude other tension-dependent effects, such as spatial separation of kinetochores from inner-centromeric Aurora B (Liu et al., 2009; Tanaka et al., 2002), or the intrinsic catch bond-like behavior of kinetochores (Akiyoshi et al., 2010). While the effect we uncovered here is specific to the kinase because we blocked phosphatase activity, it remains possible that the opposing phosphatase *in vivo* is also differentially regulated by tension. We suggest that the substrate conformation mechanism works in tandem with these other mechanisms to help ensure mitotic accuracy. Multiple force-sensitive molecules work together in many contexts, including focal adhesions (Bauer et al., 2019; del Rio et al., 2009; Kong et al., 2009; Sawada et al., 2006), adherens junctions (Buckley et al., 2014; Yao et al., 2014), muscle sarcomeres (Baumann et al., 2017; Lange et al., 2005; Puchner et al., 2008), hemostasis (Fu et al., 2017; Rognoni et al., 2012), and bacterial fimbriae (Forero et al., 2006; Le Trong et al., 2010; Thomas et al., 2002). Likewise, a multiplicity of force-sensitive elements probably also underlies the correction of erroneous kinetochore-microtubule attachments during mitosis.

## Acknowledgements

We thank Arshad Desai for antibodies. We also thank Trisha Davis, Sharona Gordon, Harmit Malik, Bill Zagotta and members of the S.B. and C.L.A. labs for critical reading of the manuscript. A.K.D. was supported by postdoctoral fellowship PF-15-139-01-CCG from the American Cancer Society. C. L. A. was supported by a Packard Fellowship 2006-30521 and NIH grants R01GM079373 and R35GM134842. S. B. was supported by NIH R01GM064386 and is also an investigator of the Howard Hughes Medical Institute.

## Author Contributions

A.K.D. conceptually designed and performed experiments, analyzed data, and wrote the initial draft of the manuscript; C.J.C. performed experiments, analyzed data and edited the manuscript. C.L.A. and S.B. conceptually designed experiments, analyzed data, and wrote the final manuscript with editorial input from A.K.D.

## Methods

### Protein preparation

The AurB* gene was created in several steps. First, the non-expressed linker between the *SLI15* and *IPL1* genes was removed on two polycistronic vectors expressing the four members of the CPC: pDD2396 (wild-type kinase) and pDD2399 (kinase-dead) (Cormier et al., 2013). A TEV cleavage site was added between the 6xHis tag and the *IPL1* gene in each plasmid and then the entire *SLI15-IPL1-TEV-6xHis* genes from these constructs were amplified via PCR with BamHI and XhoI sites on the 5’ and 3’ ends, respectively, and inserted into cut pET21b vectors (EMD Biosciences). Each kinase was then expressed as a single polypeptide with a T7 tag on its N-terminus and a 6xHis tag on its C-terminus (plasmids pSB2540 and pSB2894 for wild-type and kinase-dead, respectively). For protein preparation, each plasmid was co-transformed into the Tuner pLys BL21 (DE3) *E. coli* strain (Novagen) with a plasmid that encodes lambda phosphatase (Brading et al., 2012). A single transformant was grown to OD 0.9 and expression of both proteins induced with 500 μM IPTG. Protein was expressed for four hours at 30 °C before harvesting. Cells were resuspended in prep buffer (50 mM Tris-HCl pH 7.4, 300 mM KCl, 5 mM MgCl_2_, 5 mM ATP, 1 mM DTT, 20 mM imidazole, 10% glycerol) supplemented with 1% Triton X-100, 1 μL Turbo nuclease (Accelagen) and one tablet of Complete EDTA-free protease inhibitor (Roche) (Cormier et al., 2013) and then lysed via incubation with 0.5 mg/mL lysozyme followed by sonication. Lysate was cleared by centrifugation and then applied to a column of Ni-NTA resin (Qiagen). The column was washed with 10 column volumes (CVs) of prep buffer and then protein-bound resin was resuspended in 3 CVs of prep buffer. DTT was added to 1 mM and 3 mg of TEV protease was added for three hours of room temperature incubation. TEV protease was expressed and purified in house (Tropea et al., 2009). The resin slurry was re-applied to the empty column and the flow-through was collected along with the flow-through from one wash. This was concentrated and applied to a Superdex 200 FPLC column with storage buffer (50 mM Tris-HCl pH 7.4, 300 mM NaCl, 10% glycerol, 50 mM arginine, 50 mM glutamate). Fractions were analyzed by SDS-PAGE and those from the main peak that eluted at 72 mL were pooled and concentrated.

Kinetochores were purified from budding yeast as previously described (Akiyoshi et al., 2010). Kinetochore strains have a Dsn1 protein with 6xHis and 3xFlag tags on its C-terminus. Wild-type kinetochores were purified from strain SBY8253; Mps1-1 kinetochores were isolated from SBY8726; Ndc80-7A kinetochores were isolated from SBY8522. Yeast strains were grown asynchronously to OD 4 at room temperature, harvested, washed, and resuspended in Buffer H (25 mM HEPES pH 8.0, 150 mM KCl, 2 mM MgCl_2_, 0.1 mM EDTA pH 8.0, 0.1% NP-40, 15% glycerol) supplemented with protease inhibitors, phosphatase inhibitors and 2 mM DTT. SBY8726 for the purification of Mps1-1 kinetochores was shifted to 37 °C for the last two hours of growth. After harvest, yeast drops were frozen in liquid nitrogen and then lysed using a Freezer Mill (SPEX). Lysate was clarified via ultracentrifugation at 24,000 rpm for 90 minutes and protein-containing layer was extracted with a syringe. Extract was incubated with magnetic Dynabeads previously conjugated to anti-flag antibodies for two hours at 4°C and then washed five times in Buffer H. For optical trapping assays, kinetochores were eluted by incubating with 0.83 mg/mL 3xFlag peptide and quantified by comparing the Dsn1 silver-stained band to BSA standards. For enzyme assays, kinetochores were washed but then left attached to beads.

Dam1 complex (Dam1c) was purified via a flag-tagged Dad1 protein from SBY12464 using a similar protocol (Cheeseman et al., 2002) except the initial immunoprecipitate was first washed three times with a Buffer H with 400 mM KCl before being washed twice in regular Buffer H. The complex was eluted by incubating with 0.83 mg/mL 3xFlag peptide and quantified by comparing the Spc34 silver-stained band to BSA standards.

Taxol-stabilized microtubules were grown by incubation of a mix of unlabeled and Alexa 647-labeled purified bovine tubulin in 1xBRB80 with 4.4% DMSO, 3.2 mM MgCl_2_, and 0.8 mM GTP at 37°C for 30 minutes. Microtubules and unpolymerized tubulin were separated by ultracentrifugation and sedimented material was resuspended in 1xBRB80 with 10 μM taxol. Stabilized microtubules were used up to three days after polymerization.

### Enzyme assays

For the radioactive kinase assays, kinetochores were purified and immobilized on magnetic beads and then incubated with AurB*, AurB*-KD, or buffer with 0.2 μM hot (^32^P-γ-labeled) and 200 μM cold ATP for the indicated periods of time at room temperature. Experiments were quenched in sample buffer and separated by SDS-PAGE before being silver stained. The gel was dried and exposed to a storage phosphor screen for longer than 48 hours for detection of incorporated ^32^P.

For the bulk microtubule-binding assays, taxol-stabilized microtubules were polymerized from bovine tubulin for 20 minutes at 37°C in the presence of 0.8 mM GTP, 4.4% DMSO, 0.8xBRB80, and 3.2 mM MgCl_2_. Polymerized microtubules were sheared using 10 pumps with a syringe and 27G needle before being sedimented via ultracentrifugation at 24,000 rpm for 10 minutes. The supernatant was then removed and the pellet was resuspended in warm 1xBRB80 with 10 μM taxol. For fluorescently-labeled microtubules, eight parts bovine tubulin (10 mg/mL) was mixed with one part Alexa 648-labled bovine tubulin (~5 mg/mL) for the initial incubation.

For the microtubule binding assays shown in Figs. 2 and S2, the indicated kinetochores were purified and immobilized on beads as described above. For the experiment in Fig. 2c, Mps1-1 kinetochores were then resuspended in 1xBRB80 with 1.8 mg/mL κ-casein, 20 μM taxol, and taxol-stabilized microtubules with or without 500 μM ATP as indicated. After ten minutes, either the reaction was washed with 1xBRB80 with 20 μM taxol (BTAX2) and quenched by resuspension in 1x sample buffer (second lane), or 500 μM ATP (central lane) or buffer (other lanes) was added (1 μL into 20 μL reaction). After 20 minutes from initial resuspension, all experiments were washed with BTAX2 and quenched. Components were separated with SDS-PAGE and assayed with immunoblot. For the microtubule binding assays shown in Fig. S2, wild-type or Ndc80-7A kinetochores were incubated for five minutes in 1xBRB80 with 1.8 mg/mL κ-casein, 20 μM taxol, 500 μM ATP, and taxol-stabilized fluorescently-labeled microtubules. At *t* = 0, 1 μM AurB* or buffer was added (1.1 μL into 60 μL reaction). We used fluorescent microtubules so we could precisely quantify microtubules retained on the bead-bound kinetochores by scanning the gel for fluorescence. Timepoints were removed at 10 min and 20 minutes, washed with BTAX2 and resuspended in 1x sample buffer. Components were separated by SDS-PAGE, fluorescence scanned to visualize tubulin, and transferred to a membrane for immunoblotting to visualize Dsn1. P values were calculated by comparing means with a two-tailed unpaired t-test with Welch’s correction.

### Immunoblotting

For immunoblots, proteins were transferred from SDS-PAGE gels onto 0.22 μM cellulose paper, blocked at room temperature with 4% milk in PBST, and incubated overnight at 4 °C in primary antibody. Antibody origins and dilutions in PBST were as follows: Ndc80 N-terminus: Desai lab 1:10,000; Flag: Sigma M2 F1804 1:3000; Myc: Covance 9E10 MMS-150R 1:10,000; tubulin: EMD Millipore YL1/2 MAB1864 1:1000. Blots were then washed again with PBST and incubated with secondary antibody at room temperature. Secondary antibodies were α-mouse, α-rabbit, or α-rat horseradish peroxidase-conjugated, from GE Healthcare, and used at 1:1000 dilution in 4% milk in PBST. Blots were then washed again with PBST and ECL substrate from Thermo Scientific used to visualize the proteins.

### Optical trapping

The optical trapping was performed as described previously (Akiyoshi et al., 2010; Franck et al., 2010; Sarangapani et al., 2013), with some modifications for the flow experiments as described below. Bead conjugation and microtubule seed preparation were as previously described. Polystyrene beads (560 nm average diameter for high density experiments and 440 nm average diameter for low density experiments) conjugated to anti-His antibodies were incubated with Mps1-1 kinetochores. For the experiments conducted at high density, we incubated the beads at a concentration of 9.0 nM Dsn1 for 1 hour at 4 °C, which is >10-fold higher concentration than in our previous force-clamping experiments and potentially resulted in multiple kinetochore particles binding to a single microtubule. For the experiments conducted at low density, we incubated beads (440nm average diameter) conjugated to anti-His antibodies at a final concentration of 4.0 pM with Mps1-1 kinetochores at a final concentration of 3 nM Dsn1 for 1 hour at 4 °C. Reaction chambers were constructed on glass slides using sodium hydroxide- and ethanol-cleaned coverslips for an approximately 14 μL volume chamber. Coverslip-anchored microtubules were grown via successive introduction of: 15 μL of 10 mg/mL biotinylated bovine serum albumin, 50 μL of BRB80 (80mM K-PIPES, 1mM MgCl2, 1mM EGTA), 25 μL of 1mg/mL avidin DN, 50 μL of BRB80, 50 μL of GMPCPP-stabilized microtubule seeds, 100 μL of GTP- and κ-casein-containing BRB80, and then 50 μL of a final reaction buffer consisting of BRB80, 0.3 mg/mL κ-casein, 1 mM GTP, 0.8 mM DTT, 48 mM glucose, 1.6 mg/mL glucose oxidate, 0.3 mg/mL catalase, 3 nM purified Dam1c, 400 μM ATP, bead-bound kinetochores, and 2.6 mg/mL unpolymerized tubulin. This is similar to previous versions of the optical trap assay except with lower casein and with the addition of ATP. For these experiments, free beads were attached to coverslip bound microtubules using a lower trap stiffness of either 0.028 pN/nm for experiments conducted at 1.5 pN or 0.055 pN/nm for experiments conducted at 5 pN. Beads were dragged to the dynamic microtubule tip under a preload force of 1 pN. Once at the tip, the force was increased to either 1.5 pN or 5 pN. After one minute of tip tracking at either high or low force, 20uL of flow buffer was introduced via capillary action. Flow buffer contained BRB80, 0.3 mg/mL κ-casein, 1 mM GTP, 0.8 mM DTT, 48 mM glucose, 1.6 mg/mL glucose oxidate, 0.3 mg/mL catalase, 500 μM ATP, 0.5 μM AurB* or AurB*-KD. Flow in the chamber occurred immediately and was observed in the DIC view on the microscope. Events were observed without interference until either a rupture of the attachment occurred, the event was ended by a second bead or large piece of debris falling into the trap, the bead sticking to the coverslip, or by the bead tracking permanently back to the microtubule seed. Detachment rates were calculated by dividing the total number of detachments observed under a given condition by the total attachment time recorded. To avoid biasing the data towards higher detachment rates, we did not discard events that were interrupted and instead calculated overall detachment rates by dividing the number of detachments observed for a given condition by the total observation time after kinase flow was initiated. In all cases that describe error as “due to Poisson statistics,” uncertainties were estimated by dividing the detachment rate by the square root of the number of detachments. Custom software written in Labview (National Instruments) was used for trap instrument control and data collection. Data analysis was performed using custom software written in Igor Pro (Wavemetrics). The rupture force assay for Fig. S3a was performed under conditions identical to those previously described (Miller et al., 2016).

### Yeast Strains

Strains used in this study were constructed by standard techniques and are derivatives of W303 (*MATa ura3-1 leu2-3,112 his3-11 trp1-1 can1-100 ade2-1 bar1-1*). SBY8253 contains *DSN1-6His-3Flαg:URA3*, SBY8522 contains *DSN1-6His-3Flag:URA3 ndc80::NAT:ndc80-7A:TRP1*, SBY8726 contains *DSN1-6His-3Flag:URA3 mps1-1* and SBY12464 contains *Dad1-3Flag:TRP1*.

---

**Figure S1.**
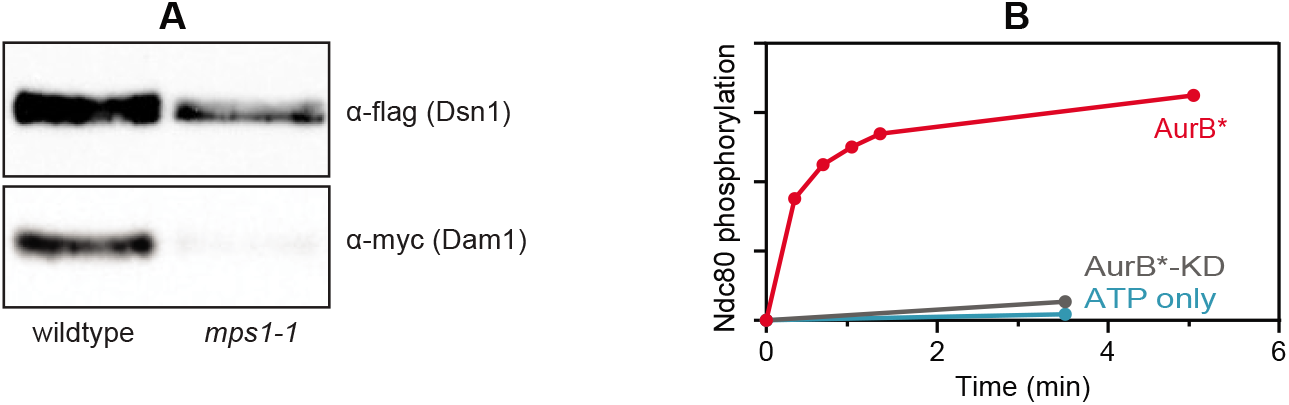
Control data related to Figure 1. (**A**) Immunoblot of purified kinetochores from either wild-type (SBY8253) or *mps1-1* cells (SBY8726) showing that Mps1-1 kinetochores lack Dam1c. (**B**) Quantification of 32P incorporation into Ndc80, based on the autoradiogram shown in Figure 1B. The red points (AurB*) are from the time course, which is shown in lanes three through seven of the gel in Figure 1B. The blue point (ATP only) is from a 3.5-min mock treatment, shown in the second to last lane of the gel in Figure 1B. The grey point (AurB*-KD) is from a 3.5-min treatment with kinase-dead mutant, shown in the last lane of the gel in Figure 1B. Units are arbitrary.

**Figure S2.**
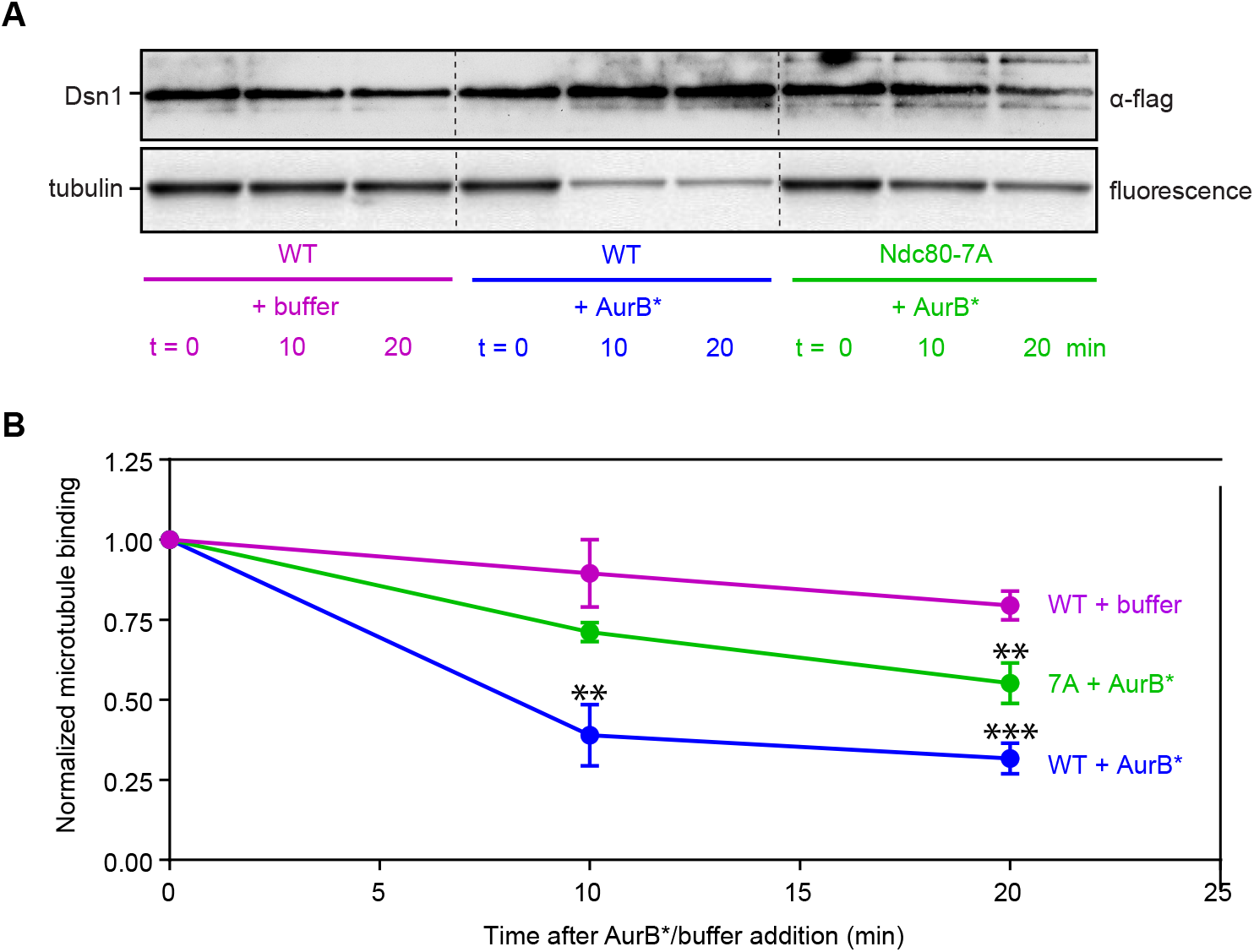
Phosphorylation of physiologically relevant sites accelerates release of microtubules from kinetochores. (**A**) Purified wild-type (WT) (SBY8253) or phospho-deficient mutant Ndc80-7A (SBY8522) kinetochores were incubated for five minutes with taxol-stabilized fluorescently-labeled microtubules in the presence of ATP. Buffer or 1 μM AurB* was added at t = 0 min and samples were removed, washed and quenched at the indicated times. Components retained after the wash were separated via SDS-PAGE and visualized by immunoblot (Dsn1-flag) and fluorescence scan (tubulin). (**B**) Bound microtubules (tubulin fluorescence) were quantified and graphed. Points are averages of three independent experiments and error bars represent standard deviation. Values were normalized to bound microtubules at t = 0. P-values between the t = 20 min timepoints using an unpaired t-test are as follows: 0.0002 (WT+buffer versus WT+AurB*), 0.0077 (WT+buffer versus 7A+AurB*), 0.0084 (WT+AurB* versus 7A+AurB*).

**Figure S3.**
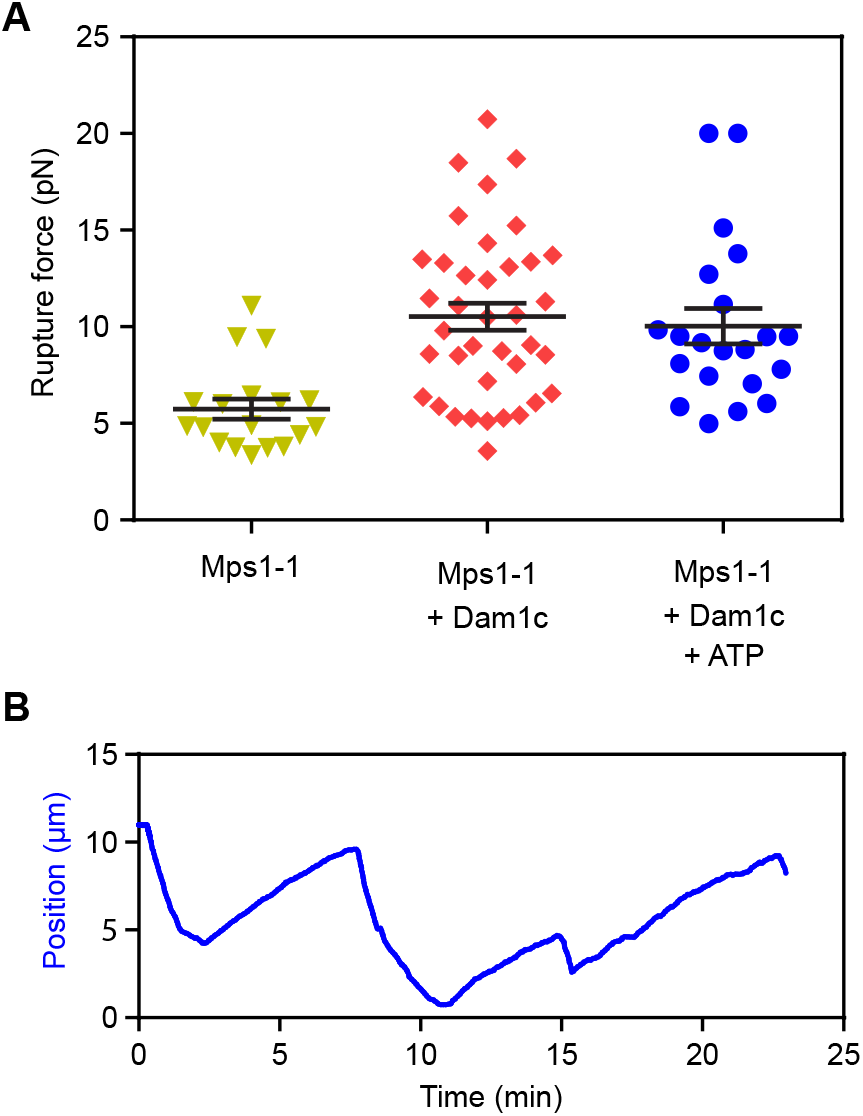
Adding Dam1c strengthens Mps1-1 kinetochores. (**A**) Rupture strengths measured for Mps1-1 kinetochores using an optical force-ramp. Where indicated, exogenously purified Dam1 complex (Dam1c) and/or ATP were included in the trapping buffer. (**B**) Trace of optical trap data for an individual force-clamp event, recorded using a bead decorated with Mps1-1 kinetochores in the presence of free purified Dam1c. The relative position versus time is plotted for a single kinetochore-decorated bead as it continuously supports 5 pN of tension while tracking with the dynamic tip of an individual microtubule during tip growth and shortening. These data (A, B) were collected using a high density of kinetochores on the trapping beads (Dsn1:bead ratio, 3,300).

**Figure S4.**
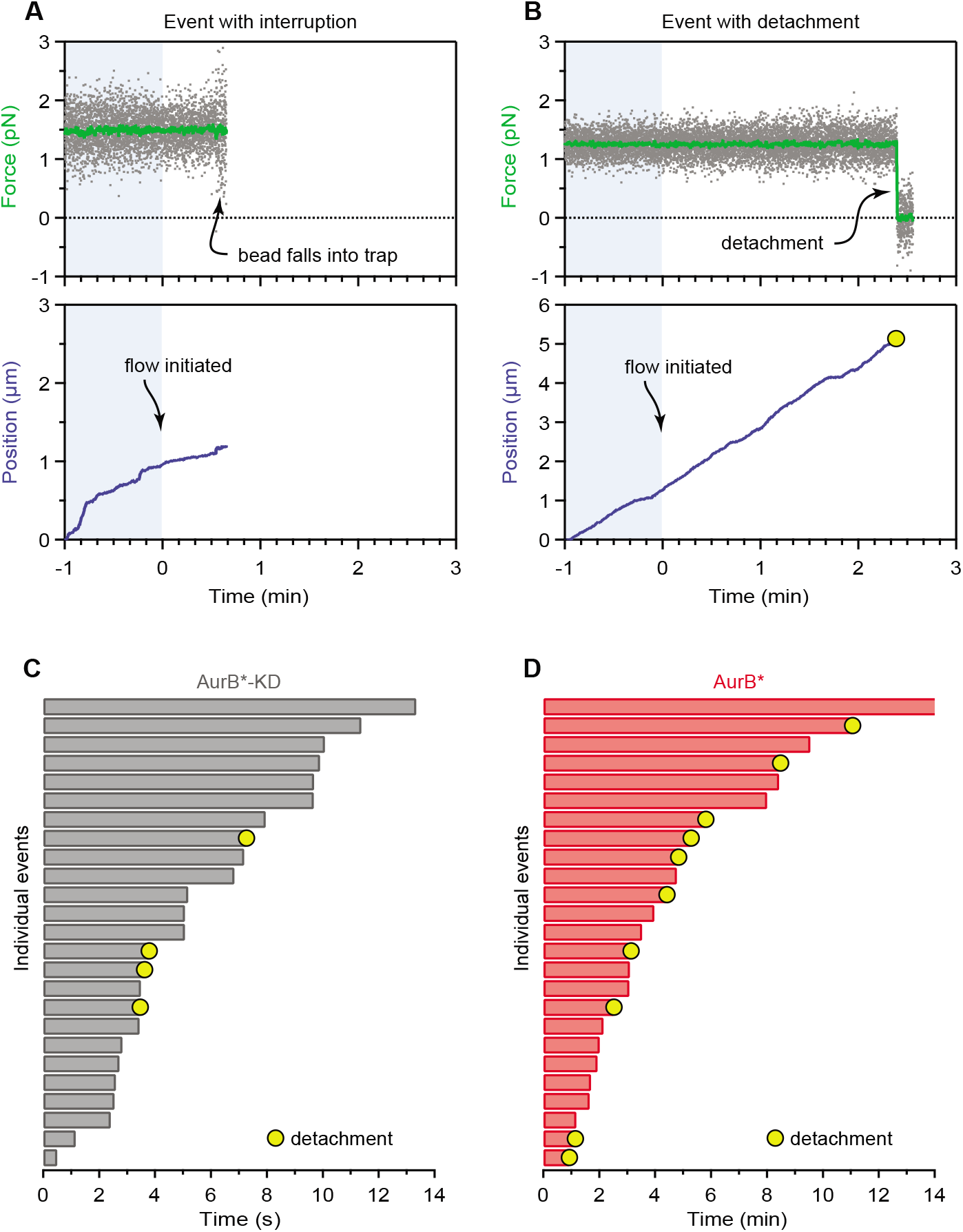
Analysis of individual events recorded using the optical trap flow assay. (**A**, **B**) Example records of optical trap data for two individual events. Gray points show the force applied and green traces show the same data after smoothing. Blue traces show relative positions of the kinetochore-decorated beads over time. Kinase introduction occurred at time 0; blue shading indicates data recorded before kinase introduction. (A) shows an event that ended when a second bead became caught in the laser trap. (B) shows an event that ended in a detachment. The yellow circle indicates when a detachment occurred. (**C**, **D**) Durations of twenty-five randomly selected kinetochore-microtubule tip attachments supporting ~1 pN of tension in the presence of either AurB*-KD (C) or AurB* (D). Events that ended in detachment are marked with yellow circles. Events that were interrupted are unmarked. Bar color indicates kinase concentration (grey, 5 μM AurB*-KD; red, 0.5 μM AurB*). These data (A - D) were collected using a high density of kinetochores on the trapping beads (Dsn1:bead ratio, 3,300).

**Figure S5.**
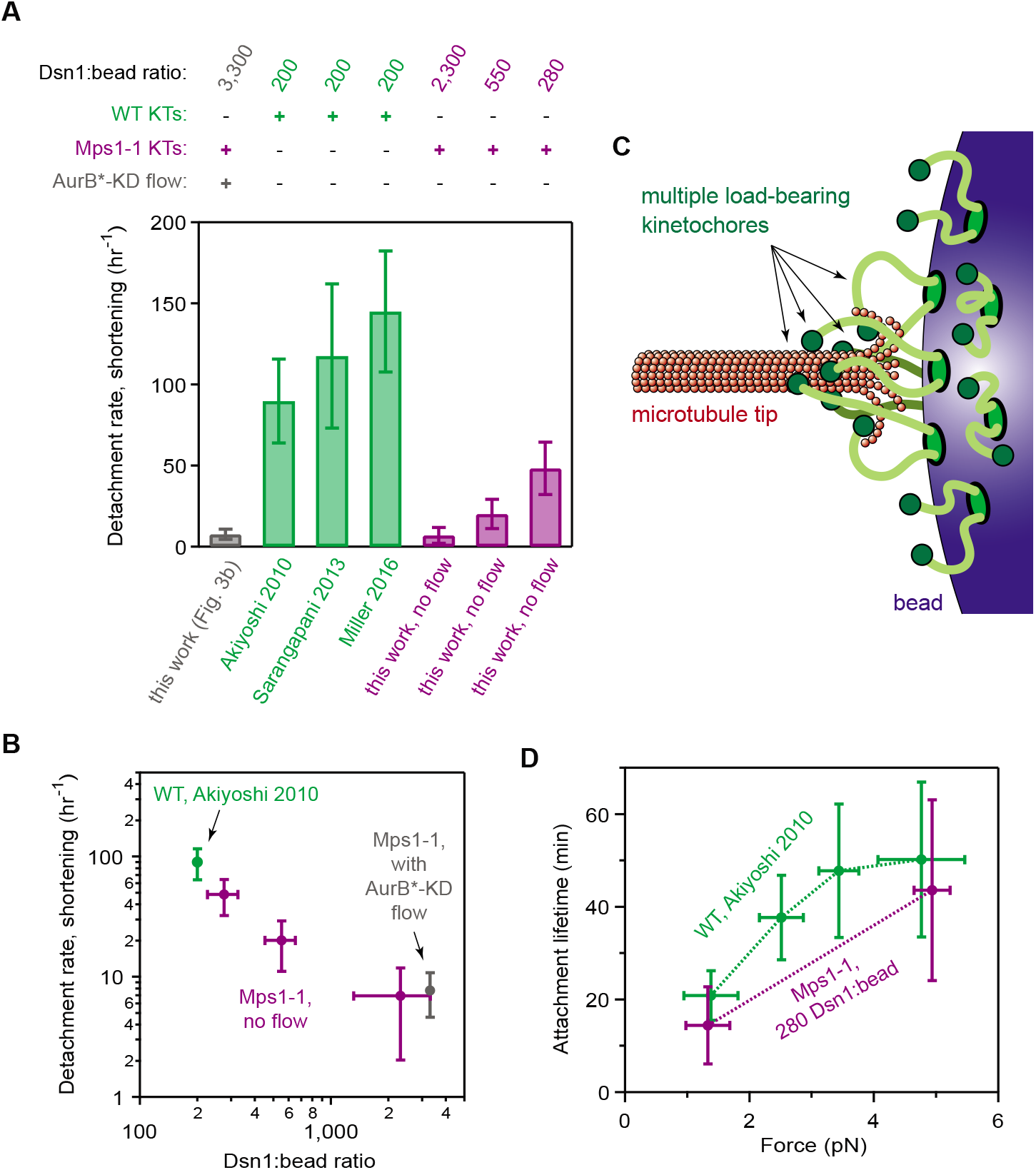
A high density of kinetochores on beads suppresses detachment, suggesting applied loads are shared by multiple kinetochores. (**A**, **B**) Detachments from shortening microtubule tips were unusually rare during initial flow assay experiments, which used a high density of Mps1-1 kinetochores on the trapping beads (AurB*-KD, grey bar; Dsn1:bead ratio, 3,300) relative to previous measurements that used beads decorated much more sparsely with wild-type kinetochores (green bars; Dsn1:bead ratio, 200; from [Akiyoshi 2010; Sarangapani 2013; Miller 2016]). Reducing the density of Mps1-1 kinetochores on the beads restored the higher detachment rates (purple bars; Dsn:bead ratios indicated). Error bars represent uncertainty due to Poisson statistics. (**C**) Schematic illustrating multiple load-bearing kinetochores attached to a single microtubule tip. If multiple kinetochores share an applied load, then the load per kinetochore can be low even when the total load is high. (**D**) With sparse decoration on the trapping beads (Dsn1:bead ratio, 280), Mps1-1 kinetochores exhibit a catch bond-like increase in attachment stability with force, consistent with previous studies using wild-type kinetochores (Dsn1:bead ratio, 200). Numbers of detachments, rate and lifetime estimates, and statistical comparisons are detailed in Supplementary Data Table 1.

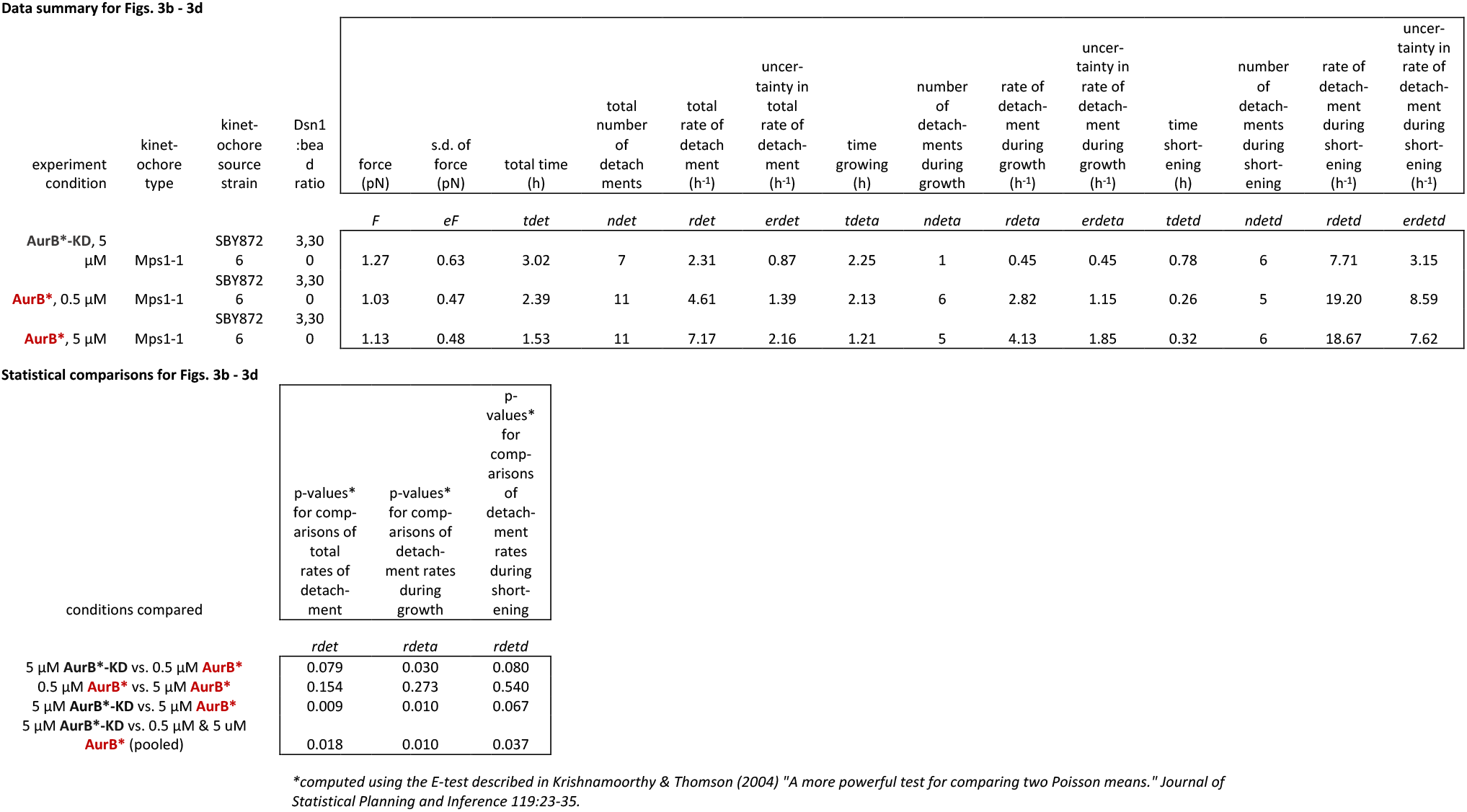

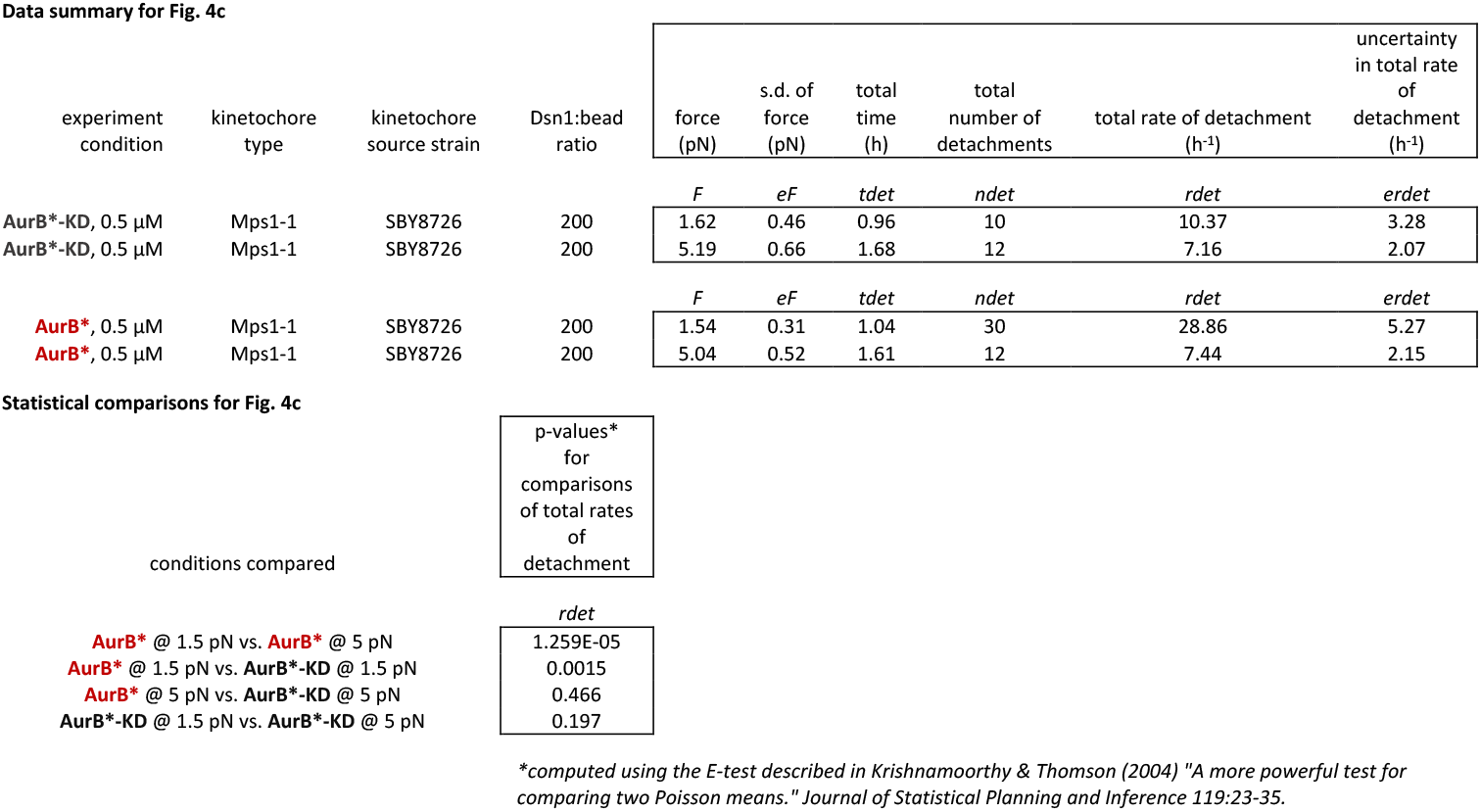

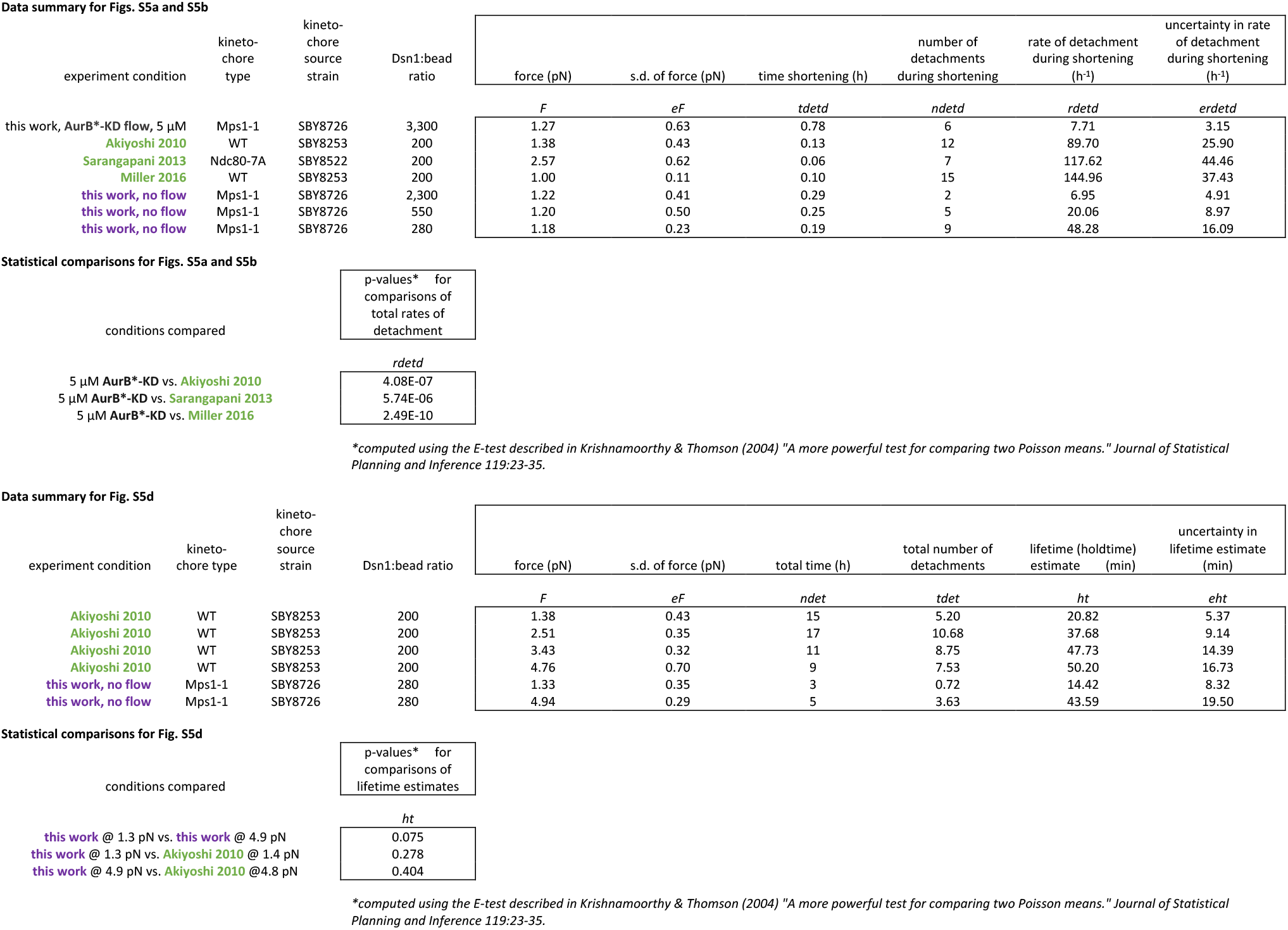

## References

Akiyoshi, B., Nelson, C.R., Ranish, J.A., and Biggins, S. (2009). Analysis of Ipl1-mediated phosphorylation of the Ndc80 kinetochore protein in *Saccharomyces cerevisiae*. Genetics 183, 1591–1595.

Akiyoshi, B., Sarangapani, K.K., Powers, A.F., Nelson, C.R., Reichow, S.L., Arellano-Santoyo, H., Gonen, T., Ranish, J.A., Asbury, C.L., and Biggins, S. (2010). Tension directly stabilizes reconstituted kinetochore-microtubule attachments. Nature 468, 576–579.

Aravamudhan, P., Felzer-Kim, I., Gurunathan, K., and Joglekar, A.P. (2014). Assembling the protein architecture of the budding yeast kinetochore-microtubule attachment using FRET. Curr Biol 24, 1437–1446.

Bauer, M.S., Baumann, F., Daday, C., Redondo, P., Durner, E., Jobst, M.A., Milles, L.F., Mercadante, D., Pippig, D.A., Gaub, H.E., et al. (2019). Structural and mechanistic insights into mechanoactivation of focal adhesion kinase. P Natl Acad Sci USA 116, 6766–6774.

Baumann, F., Bauer, M.S., Rees, M., Alexandrovich, A., Gautel, M., Pippig, D.A., and Gaub, H.E. (2017). Increasing evidence of mechanical force as a functional regulator in smooth muscle myosin light chain kinase. Elife 6.

Biggins, S., Severin, F.F., Bhalla, N., Sassoon, I., Hyman, A.A., and Murray, A.W. (1999). The conserved protein kinase Ipl1 regulates microtubule binding to kinetochores in budding yeast. Genes Dev 13, 532–544.

Brading, R.L., Abbott, W.M., Green, I., Davies, A., and McCall, E.J. (2012). Co-expression of protein phosphatases in insect cells affects phosphorylation status and expression levels of proteins. Protein Expr Purif 83, 217–225.

Broad, A.J., and DeLuca, J.G. (2020). The right place at the right time: Aurora B kinase localization to centromeres and kinetochores. Essays Biochem 64, 299–311.

Broad, A.J., DeLuca, K.F., and DeLuca, J.G. (2020). Aurora B kinase is recruited to multiple discrete kinetochore and centromere regions in human cells. J Cell Biol 219.

Buckley, C.D., Tan, J., Anderson, K.L., Hanein, D., Volkmann, N., Weis, W.I., Nelson, W.J., and Dunn, A.R. (2014). Cell adhesion. The minimal cadherin-catenin complex binds to actin filaments under force. Science 346, 1254211.

Campbell, C.S., and Desai, A. (2013). Tension sensing by Aurora B kinase is independent of survivin-based centromere localization. Nature 497, 118–121.

Chacon, J.M., Mukherjee, S., Schuster, B.M., Clarke, D.J., and Gardner, M.K. (2014). Pericentromere tension is self-regulated by spindle structure in metaphase. Journal of Cell Biology 205, 313–324.

Cheeseman, I.M., Anderson, S., Jwa, M., Green, E.M., Kang, J., Yates, J.R., Chan, C.S., Drubin, D.G., and Barnes, G. (2002). Phospho-Regulation of Kinetochore-Microtubule Attachments by the Aurora Kinase Ipl1p. Cell 111, 163–172.

Cheeseman, I.M., Chappie, J.S., Wilson-Kubalek, E.M., and Desai, A. (2006). The conserved KMN network constitutes the core microtubule-binding site of the kinetochore. Cell 127, 983–997.

Cimini, D., Wan, X., Hirel, C.B., and Salmon, E.D. (2006). Aurora kinase promotes turnover of kinetochore microtubules to reduce chromosome segregation errors. Curr Biol 16, 1711–1718.

Cormier, A., Drubin, D.G., and Barnes, G. (2013). Phosphorylation regulates kinase and microtubule binding activities of the budding yeast chromosomal passenger complex in vitro. J Biol Chem 288, 23203–23211.

del Rio, A., Perez-Jimenez, R., Liu, R., Roca-Cusachs, P., Fernandez, J.M., and Sheetz, M.P. (2009). Stretching single talin rod molecules activates vinculin binding. Science 323, 638–641.

DeLuca, J.G., Gall, W.E., Ciferri, C., Cimini, D., Musacchio, A., and Salmon, E.D. (2006). Kinetochore microtubule dynamics and attachment stability are regulated by Hec1. Cell 127, 969–982.

Dewar, H., Tanaka, K., Nasmyth, K., and Tanaka, T.U. (2004). Tension between two kinetochores suffices for their bi-orientation on the mitotic spindle. Nature 428, 93–97.

Dumont, S., Salmon, E.D., and Mitchison, T.J. (2012). Deformations within moving kinetochores reveal different sites of active and passive force generation. Science 337, 355–358.

Fischbock-Halwachs, J., Singh, S., Potocnjak, M., Hagemann, G., Solis-Mezarino, V., Woike, S., Ghodgaonkar-Steger, M., Weissmann, F., Gallego, L.D., Rojas, J., et al. (2019). The COMA complex interacts with Cse4 and positions Sli15/Ipl1 at the budding yeast inner kinetochore. Elife 8.

Foley, E.A., Maldonado, M., and Kapoor, T.M. (2011). Formation of stable attachments between kinetochores and microtubules depends on the B56-PP2A phosphatase. Nat Cell Biol 13, 1265–1271.

Forero, M., Yakovenko, O., Sokurenko, E.V., Thomas, W.E., and Vogel, V. (2006). Uncoiling mechanics of Escherichia coli type I fimbriae are optimized for catch bonds. PLoS Biol 4, e298.

Franck, A.D., Powers, A.F., Gestaut, D.R., Davis, T.N., and Asbury, C.L. (2010). Direct physical study of kinetochore-microtubule interactions by reconstitution and interrogation with an optical force clamp. Methods 51, 242–250.

Fu, H., Jiang, Y., Yang, D., Scheiflinger, F., Wong, W.P., and Springer, T.A. (2017). Flow-induced elongation of von Willebrand factor precedes tension-dependent activation. Nat Commun 8, 324.

Gutierrez, A., Kim, J.O., Umbreit, N.T., Asbury, C.L., Davis, T.N., Miller, M.P., and Biggins, S. (2020). Cdk1 Phosphorylation of the Dam1 Complex Strengthens Kinetochore-Microtubule Attachments. Curr Biol 30, 4491–4499 e4495.

Hengeveld, R.C.C., Vromans, M.J.M., Vleugel, M., Hadders, M.A., and Lens, S.M.A. (2017). Inner centromere localization of the CPC maintains centromere cohesion and allows mitotic checkpoint silencing. Nat Commun 8, 15542.

Joglekar, A.P., Bloom, K., and Salmon, E.D. (2009). In vivo protein architecture of the eukaryotic kinetochore with nanometer scale accuracy. Curr Biol 19, 694–699.

Kong, F., Garcia, A.J., Mould, A.P., Humphries, M.J., and Zhu, C. (2009). Demonstration of catch bonds between an integrin and its ligand. Journal of Cell Biology 185, 1275–1284.

Krenn, V., and Musacchio, A. (2015). The Aurora B Kinase in Chromosome Bi-Orientation and Spindle Checkpoint Signaling. Front Oncol 5, 225.

Kruse, T., Zhang, G., Larsen, M.S., Lischetti, T., Streicher, W., Kragh Nielsen, T., Bjorn, S.P., and Nilsson, J. (2013). Direct binding between BubR1 and B56-PP2A phosphatase complexes regulate mitotic progression. J Cell Sci 126, 1086–1092.

Lampson, M.A., Renduchitala, K., Khodjakov, A., and Kapoor, T.M. (2004). Correcting improper chromosome-spindle attachments during cell division. Nat Cell Biol 6, 232–237.

Lange, S., Xiang, F., Yakovenko, A., Vihola, A., Hackman, P., Rostkova, E., Kristensen, J., Brandmeier, B., Franzen, G., Hedberg, B., et al. (2005). The kinase domain of titin controls muscle gene expression and protein turnover. Science 308, 1599–1603.

Le Trong, I., Aprikian, P., Kidd, B.A., Forero-Shelton, M., Tchesnokova, V., Rajagopal, P., Rodriguez, V., Interlandi, G., Klevit, R., Vogel, V., et al. (2010). Structural basis for mechanical force regulation of the adhesin FimH via finger trap-like beta sheet twisting. Cell 141, 645–655.

Liu, D., Vader, G., Vromans, M.J., Lampson, M.A., and Lens, S.M. (2009). Sensing chromosome bi-orientation by spatial separation of aurora B kinase from kinetochore substrates. Science 323, 1350–1353.

Liu, D., Vleugel, M., Backer, C.B., Hori, T., Fukagawa, T., Cheeseman, I.M., and Lampson, M.A. (2010). Regulated targeting of protein phosphatase 1 to the outer kinetochore by KNL1 opposes Aurora B kinase. J Cell Biol 188, 809–820.

London, N., Ceto, S., Ranish, J.A., and Biggins, S. (2012). Phosphoregulation of Spc105 by Mps1 and PP1 regulates Bub1 localization to kinetochores. Curr Biol 22, 900–906.

Maresca, T.J., and Salmon, E.D. (2009). Intrakinetochore stretch is associated with changes in kinetochore phosphorylation and spindle assembly checkpoint activity. J Cell Biol 184, 373–381.

Meyer, R.E., Kim, S., Obeso, D., Straight, P.D., Winey, M., and Dawson, D.S. (2013). Mps1 and Ipl1/Aurora B act sequentially to correctly orient chromosomes on the meiotic spindle of budding yeast. Science 339, 1071–1074.

Miller, M.P., Asbury, C.L., and Biggins, S. (2016). A TOG Protein Confers Tension Sensitivity to Kinetochore-Microtubule Attachments. Cell 165, 1428–1439.

Mukherjee, S., Sandri, B.J., Tank, D., McClellan, M., Harasymiw, L.A., Yang, Q., Parker, L.L., and Gardner, M.K. (2019). A Gradient in Metaphase Tension Leads to a Scaled Cellular Response in Mitosis. Developmental Cell 49, 63-+.

Nicklas, R.B., and Koch, C.A. (1969). Chromosome micromanipulation. 3. Spindle fiber tension and the reorientation of mal-oriented chromosomes. J Cell Biol 43, 40–50.

Nicklas, R.B., and Ward, S.C. (1994). Elements of error correction in mitosis: microtubule capture, release, and tension. J Cell Biol 126, 1241–1253.

Puchner, E.M., Alexandrovich, A., Kho, A.L., Hensen, U., Schafer, L.V., Brandmeier, B., Grater, F., Grubmuller, H., Gaub, H.E., and Gautel, M. (2008). Mechanoenzymatics of titin kinase. P Natl Acad Sci USA 105, 13385–13390.

Rognoni, L., Stigler, J., Pelz, B., Ylanne, J., and Rief, M. (2012). Dynamic force sensing of filamin revealed in single-molecule experiments. Proc Natl Acad Sci U S A 109, 19679–19684.

Ruchaud, S., Carmena, M., and Earnshaw, W.C. (2007). Chromosomal passengers: conducting cell division. Nat Rev Mol Cell Biol 8, 798–812.

Sarangapani, K.K., Akiyoshi, B., Duggan, N.M., Biggins, S., and Asbury, C.L. (2013). Phosphoregulation promotes release of kinetochores from dynamic microtubules via multiple mechanisms. Proc Natl Acad Sci U S A 110, 7282–7287.

Sarangapani, K.K., Koch, L.B., Nelson, C.R., Asbury, C.L., and Biggins, S. (2021). Kinetochore-associated Mps1 regulates the strength of kinetochore-microtubule attachments via Ndc80 phosphorylation. bioRxiv.

Sawada, Y., Tamada, M., Dubin-Thaler, B.J., Cherniavskaya, O., Sakai, R., Tanaka, S., and Sheetz, M.P. (2006). Force sensing by mechanical extension of the Src family kinase substrate p130Cas. Cell 127, 1015–1026.

Smith, C.A., McAinsh, A.D., and Burroughs, N.J. (2016). Human kinetochores are swivel joints that mediate microtubule attachments. Elife 5.

Suijkerbuijk, S.J., Vleugel, M., Teixeira, A., and Kops, G.J. (2012). Integration of kinase and phosphatase activities by BUBR1 ensures formation of stable kinetochore-microtubule attachments. Dev Cell 23, 745–755.

Tanaka, T.U., Rachidi, N., Janke, C., Pereira, G., Galova, M., Schiebel, E., Stark, M.J., and Nasmyth, K. (2002). Evidence that the Ipl1-Sli15 (Aurora kinase-INCENP) complex promotes chromosome bi-orientation by altering kinetochore-spindle pole connections. Cell 108, 317–329.

Thomas, W.E., Trintchina, E., Forero, M., Vogel, V., and Sokurenko, E.V. (2002). Bacterial adhesion to target cells enhanced by shear force. Cell 109, 913–923.

Tien, J.F., Umbreit, N.T., Zelter, A., Riffle, M., Hoopmann, M.R., Johnson, R.S., Fonslow, B.R., Yates, J.R., 3rd, MacCoss, M.J., Moritz, R.L., et al. (2014). Kinetochore biorientation in Saccharomyces cerevisiae requires a tightly folded conformation of the Ndc80 complex. Genetics 198, 1483–1493.

Tropea, J.E., Cherry, S., and Waugh, D.S. (2009). Expression and purification of soluble His(6)-tagged TEV protease. Methods Mol Biol 498, 297–307.

Uchida, K.S., Takagaki, K., Kumada, K., Hirayama, Y., Noda, T., and Hirota, T. (2009). Kinetochore stretching inactivates the spindle assembly checkpoint. J Cell Biol 184, 383–390.

Uchida, K.S.K., Jo, M., Nagasaka, K., Takahashi, M., Shindo, N., Shibata, K., Tanaka, K., Masumoto, H., Fukagawa, T., and Hirota, T. (2021). Kinetochore stretching-mediated rapid silencing of the spindle-assembly checkpoint required for failsafe chromosome segregation. Curr Biol 31, 1581–1591 e1583.

Umbreit, N.T., Gestaut, D.R., Tien, J.F., Vollmar, B.S., Gonen, T., Asbury, C.L., and Davis, T.N. (2012). The Ndc80 kinetochore complex directly modulates microtubule dynamics. Proc Natl Acad Sci U S A 109, 16113–16118.

Wan, X., O’Quinn, R.P., Pierce, H.L., Joglekar, A.P., Gall, W.E., DeLuca, J.G., Carroll, C.W., Liu, S.T., Yen, T.J., McEwen, B.F., et al. (2009). Protein architecture of the human kinetochore microtubule attachment site. Cell 137, 672–684.

Willems, E., Dedobbeleer, M., Digregorio, M., Lombard, A., Lumapat, P.N., and Rogister, B. (2018). The functional diversity of Aurora kinases: a comprehensive review. Cell Div 13, 7.

Yao, M., Qiu, W., Liu, R., Efremov, A.K., Cong, P., Seddiki, R., Payre, M., Lim, C.T., Ladoux, B., Mege, R.M., et al. (2014). Force-dependent conformational switch of alpha-catenin controls vinculin binding. Nat Commun 5, 4525.

Yue, Z., Carvalho, A., Xu, Z., Yuan, X., Cardinale, S., Ribeiro, S., Lai, F., Ogawa, H., Gudmundsdottir, E., Gassmann, R., et al. (2008). Deconstructing Survivin: comprehensive genetic analysis of Survivin function by conditional knockout in a vertebrate cell line. J Cell Biol 183, 279–296.

